# Improving Proteinuria Screening with Mailed Smartphone Urinalysis Testing in Previously Unscreened Patients with Hypertension: a Randomized Controlled Trial

**DOI:** 10.1101/536995

**Authors:** Julia Leddy, Jamie A. Green, Christina Yule, Juliann Molecavage, Josef Coresh, Alex R Chang

## Abstract

**Background:** Proteinuria screening is recommended for patients with hypertension to screen for kidney disease and identify those at elevated risk for cardiovascular disease. However, screening rates among hypertensive patients are low. Home testing strategies may be useful in improving proteinuria screening adherence.

**Methods:** We conducted an individual-level, randomized trial at 55 primary care clinic sites in the Geisinger Health System to evaluate the effectiveness of a strategy using home smartphone urinalysis test (Dip.io) to complete proteinuria screening in previously unscreened non-diabetic patient portal users with hypertension. All patients received an educational letter and a standing urinalysis lab order, and then were randomized to control (usual care) or intervention. Intervention arm participants were invited to complete proteinuria screening with a mailed home smartphone urinalysis test. Co-primary outcomes were completion of proteinuria screening and number of albuminuria cases (albumin/creatinine ratio [ACR] ≥ 30 mg/g or protein/creatinine ratio ≥ 150 mg/g) at the end of 3 months. We also evaluated patient satisfaction with the home test, and compliance with recommendations for patients with newly detected albuminuria.

**Results:** A total of 999 patients were randomized to intervention or control. Out of 499 patients assigned to the intervention arm, 253 were reached by phone, and 69/97 (71.1%) consented patients completed the home test. Overall, the intervention increased proteinuria screening completion (28.9% vs. 18.0%; p<0.001) with no effect on the number of albuminuria cases (4 vs. 4) although only 6/57 (10.5%) patients with trace or 1+ urine dipstick protein had a follow-up quantitative test. Among the 55 patients who completed a survey after the home test, 89% preferred testing at home rather than the physician’s office.

**Conclusions:** A strategy using a home urinalysis smartphone test increased proteinuria screening rates in previously unscreened patients with hypertension and may be useful in increasing rates of proteinuria screening compliance. Future studies are needed to determine whether improving early detection of kidney disease can improve future kidney health.

**Trial Registration:** Clinical Trial Registry: NCT03470701

https://clinicaltrials.gov/ct2/show/NCT03470701

## Background

Presence of albuminuria or proteinuria is associated with increased risk for cardiovascular and kidney disease[1, 2]. While guidelines recommend screening for kidney disease using urinalysis dipstick for patients with hypertension (and urine albumin/creatinine ratio [ACR] for those with diabetes or chronic kidney disease [CKD])[3, 4], screening remains suboptimal[5–8]. Top research priorities raised by patients with CKD include improving strategies of early kidney disease identification, which could allow implementation of interventions to slow progression of disease [9, 10]. Presence of albuminuria or proteinuria can also be helpful in risk stratification to determine which patients may benefit from management with blood pressure medications, inhibition of the renin-angiotensin-aldosterone system, and statins[11].

One potential barrier to screening is that patients may not be able to provide a urine sample in the office. Extending testing to the home may be a novel strategy to improve proteinuria screening. For example, use of mailed fecal occult blood testing kits has been shown to double compliance with colorectal cancer screening[12, 13]. The objective of this randomized controlled trial was to evaluate the effectiveness of home urinalysis testing, using a smartphone urinalysis kit (Dip.io), among previously unscreened patients with hypertension receiving care at the Geisinger Health System.

## Methods

This study was an individual-level, randomized trial that included 999 patients at 55 primary care clinic sites in the Geisinger Health System, a large integrated health system in rural Pennsylvania. Funding was provided by the National Kidney Foundation (NKF), and the Geisinger Institutional Review Board approved the protocol (2017-0516). Health care providers and pharmacists across the system received education about management of proteinuria as well as information about the research study in small group lectures.

### Study Population

We used electronic health record data to identify eligible patients with hypertension who had never undergone proteinuria screening by any method. Inclusion criteria included Geisinger primary care patients ≥ 18 years of age with a hypertension diagnosis, blood pressure ≥ 130/80 mmHg at last outpatient visit, and active patient portal use with a listed phone number and email address. Main exclusion criteria included diabetes, end-stage renal disease, and estimated glomerular filtration rate (eGFR) < 15 ml/min/1.73 m^2^.

### Participant Flow and Intervention

A total of 999 patients met the inclusion/exclusion criteria and received a mailed reminder to complete proteinuria screening, a National Kidney Foundation educational booklet about the importance of proteinuria screening, and a standing Geisinger lab urinalysis order March 19-21, 2018. Since proteinuria screening is standard-of-care for patients with hypertension, informed consent was not required for this initial contact. A computer program was then used to randomize patients 1:1 to the intervention or control arm, stratified by baseline eGFR < 60 ml/min/1.73m^2^ since CKD is also an indication for proteinuria screening.

Patients in the control arm received no intervention, whereas patients in the intervention arm received a notification letter about the home urine smartphone test approximately 2 weeks later, with an option to opt-out of further contact by the research team. This was followed by telephone calls by the Geisinger Survey unit, who made up to 7 attempts and left up to 3 voicemails. We also conducted weekly data pulls to identify and remove any patients from the call list who had already completed proteinuria screening at the Geisinger lab. Patients who provided consent were sent a text message link to download the dip.io app from the Apple Store or Google Play (Figure 1). The testing kit, along with a project leaflet, were shipped by Healthy.io using a third-party fulfillment service. If participants had not completed the test within the next week, Healthy.io’s call center contacted participants to validate receipt of kit and application and trouble-shoot any issues that may have prevented the participants from completing the test.

**Figure 1.**
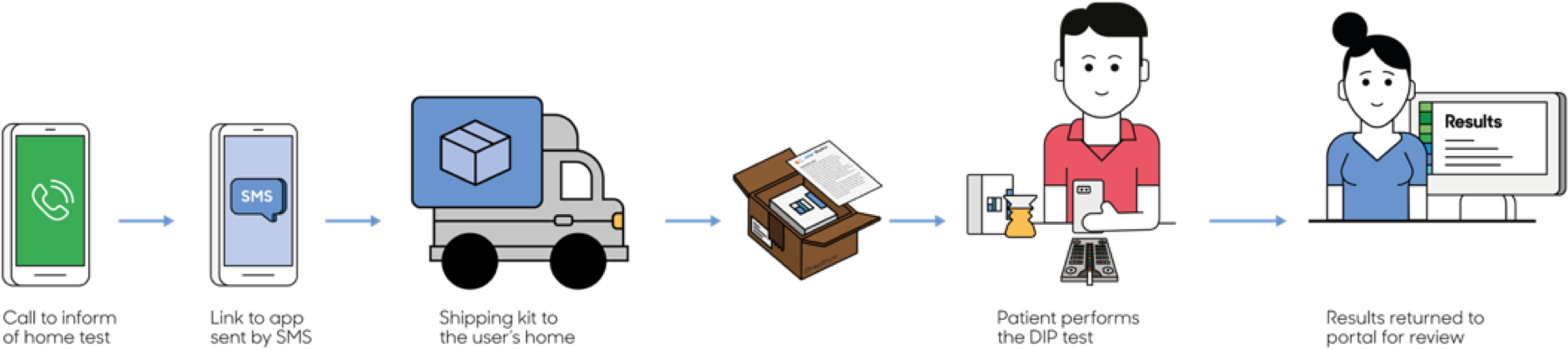
Dip.io Urinalysis Testing Process. Patients in the intervention arm who provided verbal consent were sent a text message and a link to download the dip.io app from the Apple Store or Google Play. Dip.io tests were then shipped to participants who opened the app, followed directions provided on-screen, collected urine in the provided container, dipped the urinalysis dipstick, placed the dipstick on the color board, and scanned the dipstick and color board using the app. For this project, results were transmitted to a Health Insurance Portability and Accountability Act (HIPAA)-compliant website portal, which was accessed by the research team.

The dip.io test consists of a home test kit and a smartphone application. The kit consists of a standard 10 parameter urinalysis dipstick, a custom designed urine cup and a color-board, which enables accurate analysis in different lighting environments. To conduct the test, patients open the app, follow directions provided on-screen, collect urine in the provided container, dip the urinalysis dipstick (Acon Mission), place the dipstick on the color board, and then scan the dipstick and color board using the app. For this project, results were transmitted to a Health Insurance Portability and Accountability Act (HIPAA)-compliant website portal, which was accessed by the research team. The study team contacted patients with abnormal home test results, and ordered confirmation ACR testing for trace or greater urine protein on urinalysis, or repeat urinalysis testing at a Geisinger laboratory for other urinalysis abnormalities along with notification to the PCP. For patients with detected albuminuria, the study team sent a notification to the PCP with the following guideline-based recommendations: treat to office blood pressure < 130/80 mmHg, treat with ACEI or ARB and statin. Providers had the option of referral for further management by pharmacists, who received training in management of albuminuria and hypertension. PCPs were blinded to their patients’ assignment with the exception that if an abnormal home test resulted, PCPs were later informed of the result. Final outcome data was assessed by a blinded member of the research team (JL).

**Figure 2.**
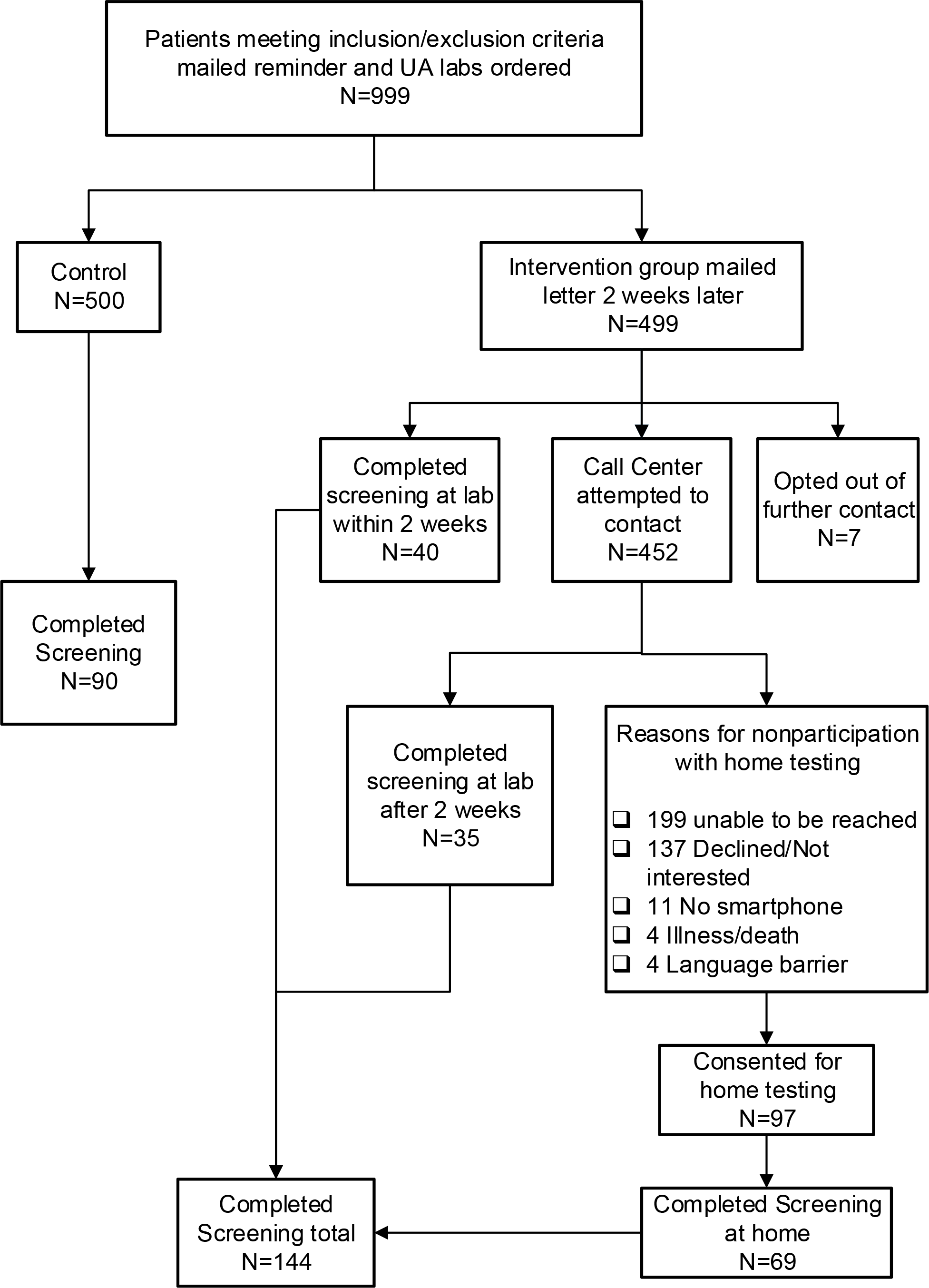
Study Flow. A total of 999 patients met inclusion/exclusion criteria and were randomized; all were included in analyses. In the control arm, 90/500 (18.0%) completed proteinuria screening tests. In the intervention arm, a total of 144/499 (28.9%) patients completed proteinuria screening tests, including 69 which were completed with the home testing kit.

### Outcomes

Co-primary outcomes were completion of proteinuria screening by any method and number of quantified albuminuria cases (ACR ≥ 30 mg/g or protein/creatinine ratio ≥ 150 mg/g) at the end of 3 months (6/20/18). An exploratory outcome examined was the number of patients with trace or 1+ urine protein tests at the end of 3 months.

### Analytic Considerations

With a sample size of 1000 total patients enrolled in the trial and assumptions of albuminuria prevalence of 20%[14], screening compliance rates of 40% in the intervention group and 10% in the control group, we estimated that we would have >90% power to detect a difference in screening compliance between groups, and >90% power to detect a difference in detected number of albuminuria cases.

We conducted intention-to-treat analyses and used logistic regression to examine the effects of the intervention on screening outcomes. Satisfaction with the home test was evaluated by a survey administered by smartphone after test completion. P values <0.05 were considered statistically significant without correction for multiple comparisons, and analyses were completed using STATA version 15.1.

## Results

Of the 999 qualifying patients in the trial, mean age was 50.5 ± 11.8 years, 40.6% were female, 93.8% were white, mean body mass index was 34.7 ± 8.5 kg/m^2^, mean systolic blood pressure was 139.8 ± 14.0 mmHg, mean diastolic blood pressure was 86.2 ± 9.2 mmHg, 70.0% were on blood pressure medications, 80.3% had eGFR ≥ 60 ml/min/1.73m^2^, 2.2% had eGFR 15-59 ml/min/1.73m^2^, and 17.5% had missing eGFR (Table 1). All participants were included in analyses.

**Table 1.**
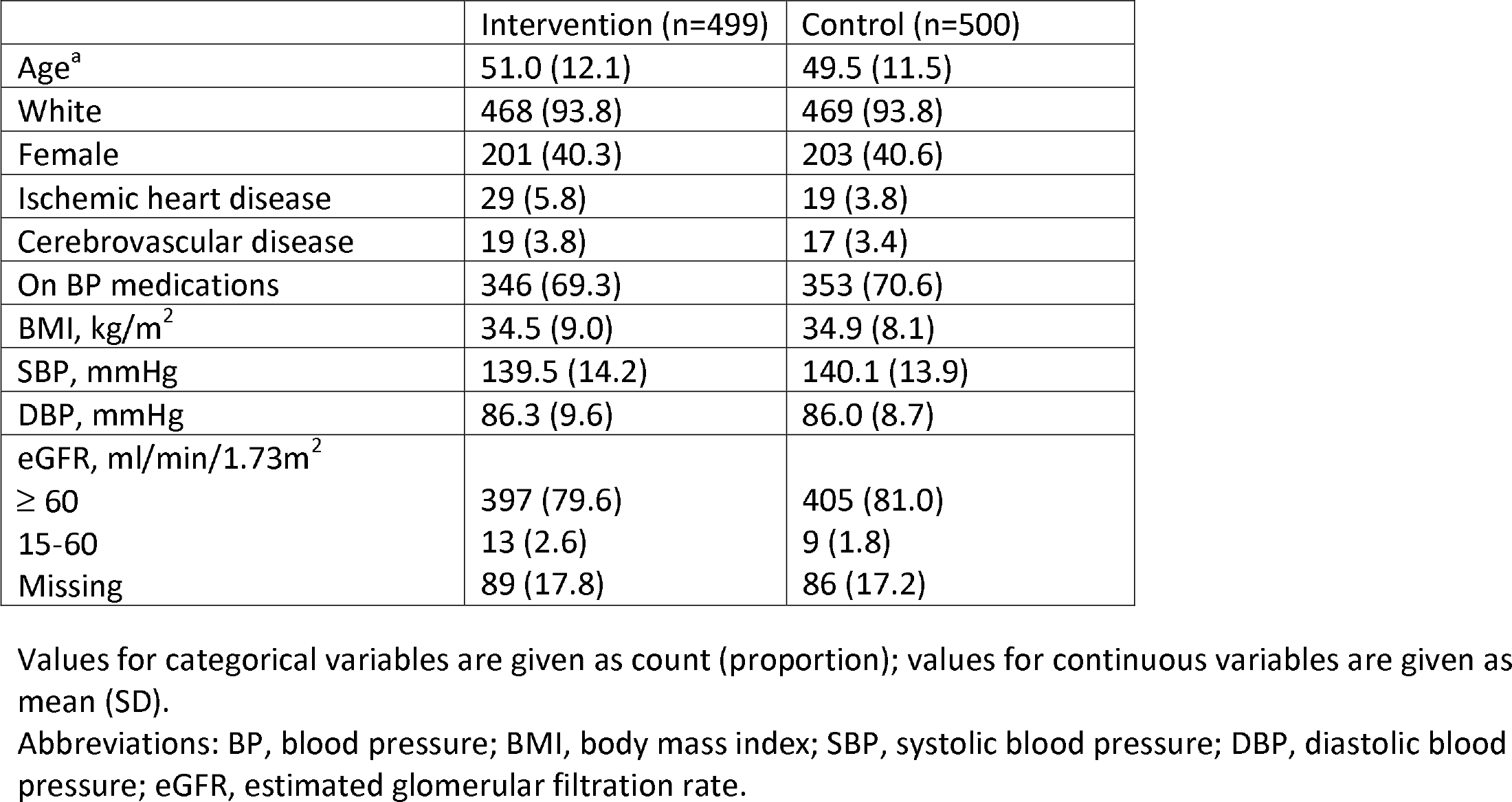
Baseline Characteristics.

### Participation in Home Testing in the Intervention Arm

Of the 499 patients in the intervention arm, 7 (1.4%) opted out of further contact after receiving the notification letter about the opportunity to use mailed smartphone urinalysis kit for proteinuria testing. Two weeks after the notification letter was sent, 40 patients in the intervention arm had already completed proteinuria screening at an outpatient Geisinger lab (Figure 1). Of the remaining 452 participants, the survey unit successfully reached 253 patients, and 35 participants completed proteinuria screening at an outpatient Geisinger lab. Reasons for declining to participate with home testing included lack of interest (137), no smartphone (11), illness/death (4), and language barrier (4). Out of 97 patients who consented for home testing, 69 (71.1%) completed the home test, with 11 patients having trace protein and 1 patient having 1+ protein.

### Effect of the Intervention on Screening Outcomes

Overall, the proportion of patients who completed proteinuria screening after 3 months was greater in the intervention arm than in the control arm (28.9% vs. 18.0%; odds ratio [OR] 1.85, 95% confidence interval [CI]: 1.37-2.49) (Table 2). There was no significant difference between the intervention and control arms in the number of detected albuminuria cases (4 vs. 4; OR 1.00, 95% CI: 0.25-4.03) or an exploratory outcome of detected trace or 1+ urine dipstick protein (34 vs. 23; OR 1.52, 95% CI: 0.88-2.61). Notably, only 3/23 (13%) patients in the control group and 3/34 (9%) patients in the intervention group with trace or 1+ protein on urine dipstick had a quantitative test (ACR or protein/creatinine ratio) checked over the 3-month study period.

**Table 2.** Effect of the Intervention on Screening Outcomes.

|  | Intervention | Control | OR (95% CI) | P value |
| --- | --- | --- | --- | --- |
| Screened by any method | 144 (28.9%) | 90 (18.0%) | 1.85 (1.37, 2.49) | <0.001 |
| Quantified albuminuria cases (ACR 30+ mg/g or PCR 150+ mg/g) | 4 (0.8%) | 4 (0.8%) | 1.00 (0.25, 4.03) | 1.00 |
| Detected trace or 1+ urine protein on dipstick | 34 (6.8%) | 23 (4.6%) | 1.52 (0.88, 2.61) | 0.13 |

Abbreviations: ACR (albumin/creatinine ratio), OR (odds ratio)

### Satisfaction with Home Testing

Surveys were completed by 55/69 (79.7% response rate) patients after using the smartphone testing kit: 98% rated ease of use as “easy” or “very easy”; 93% reported no problems with the device; 89% preferred testing at home rather than the physician’s office; mean score for whether they would recommend home urine testing to a friend or colleague was 8.9 (0-10, 10=highest; net promoter score 62[15]). No harms or unintended effects were noted by participants.

## Discussion

In this multi-site, randomized controlled trial in a large, rural, integrated health system, we found that use of mailed smartphone urinalysis kits increased rates of albuminuria screening among previously unscreened patients with hypertension. While only 19% in the intervention arm received the home testing kit, 71% of those who received the kit completed testing. Those who completed home testing were highly satisfied and preferred testing at home vs. the lab, suggesting that home testing may be a useful strategy to improve compliance with proteinuria screening.

While our intervention improved screening, there was no difference in the number of detected albuminuria cases, confirmed by quantitative ACR or PCR testing. Several factors may have limited our ability to detect more albuminuria cases. Foremost, we required informed consent to send home testing kits to patients as the study was conducted prior to the 2018 FDA approval of dip.io. Thus, only 97/499 (19.4%) in the intervention arm provided consent to receive the home testing kit. Conducting this trial with a full waiver of consent would have allowed more patients in the intervention arm to receive the home testing kits and likely more screening completions. Second, we targeted non-diabetic, hypertensive patients, who had never received screening. This resulted in a study population that was at the healthier spectrum of at-risk patients who should be screened for albuminuria/proteinuria. Lastly, very few patients with trace or 1+ protein underwent confirmation testing by quantitative ACR or protein/creatinine ratio, over a relatively short follow-up period. It is possible that traveling to the laboratory for confirmation testing may pose an additional barrier; future studies should explore confirming albuminuria using home testing kits.

Other studies have described similarly suboptimal rates of screening among patients at risk of CKD and even patients with eGFR < 60 ml/min/1.73m^2^. For example, in a study of stage 3-4 CKD patients receiving care from 2004-2008 in a group practice in eastern Massachusetts, only 30% had annual urine protein monitoring completed[16]. Even in an optimal setting where clinical decision support in the form of a CKD checklist embedded in the EHR was implemented in a prospective study, annual ACR measurement still was only able to be improved to 73% [17]. Thus, use of home testing may have a role in further improving adherence to albuminuria/proteinuria screening.

There were several limitations in this proof-of-concept study. Due to the requirement of having a smartphone to test the urinalyses, we limited our mostly white, patient population to active patient portal users, and results may not be generalizable to non-portal users. While smartphone ownership has risen from 35% in 2011 to 77% in 2018[18], other screening strategies may also be needed to reach those without access to smartphones. Whether or not early detection of albuminuria can improve outcomes will require future large-scale studies with longer follow-up and consideration of strategies to increase patient engagement and provider adherence to clinical guidelines for albuminuria.

## Conclusions

Our intervention using smartphone-based home testing was successful in increasing proteinuria screening rates in previously unscreened patients with hypertension and may be preferable for some patients. We also found that few patients with trace or 1+ urine protein underwent follow-up ACR testing, limiting the effectiveness of a urine dipstick-first screening strategy. Further research is needed to evaluate home testing strategies using quantitative or semi-quantitative ACR for screening and confirmation of albuminuria, and to determine whether increasing albuminuria screening can improve patient-centered kidney and cardiovascular outcomes.

## Supporting information

Consort Checklist

## List of abbreviations

ACR: Albumin/creatinine ratio
CKD: chronic kidney disease
NKF: National Kidney Foundation
eGFR: estimated glomerular filtration rate
PI: principal investigator
MTM: medication therapy management
OR: odds ratio
CI: confidence interval
FDA: Federal Drug Administration

## Declarations

Ethics approval and consent to participate: The Geisinger Institutional Review Board approved the protocol (2017-0516). Informed consent was obtained verbally for patients in the intervention arm to receive the home testing kit, which was not FDA-approved at the time of the research trial. Verbal consent was obtained as written consent would have required patients to come in person, limiting generalizability. Informed consent was waived for patients in the control arm as patients in the control received routine medical care. This study adheres to CONSORT guidelines (Supplemental Materials).

### Consent for publication

Not applicable.

### Availability of data and materials

The datasets generated during this study are not publicly available, but are available from the corresponding author on reasonable request.

### Authors’ contributions

Research idea and study design: ARC, CY, JAG, JM, JC; data acquisition: CY, JM, CY; data analysis/interpretation/statistical analysis: JL, ARC; supervision or mentorship: ARC. All authors read and approved the final manuscript.

## Acknowledgments

Not applicable

