## Supplementary material for "Improving Proteinuria Screening with Mailed Smartphone Urinalysis Testing in Previously Unscreened Patients with Hypertension: a Randomized Controlled Trial": Consort Checklist

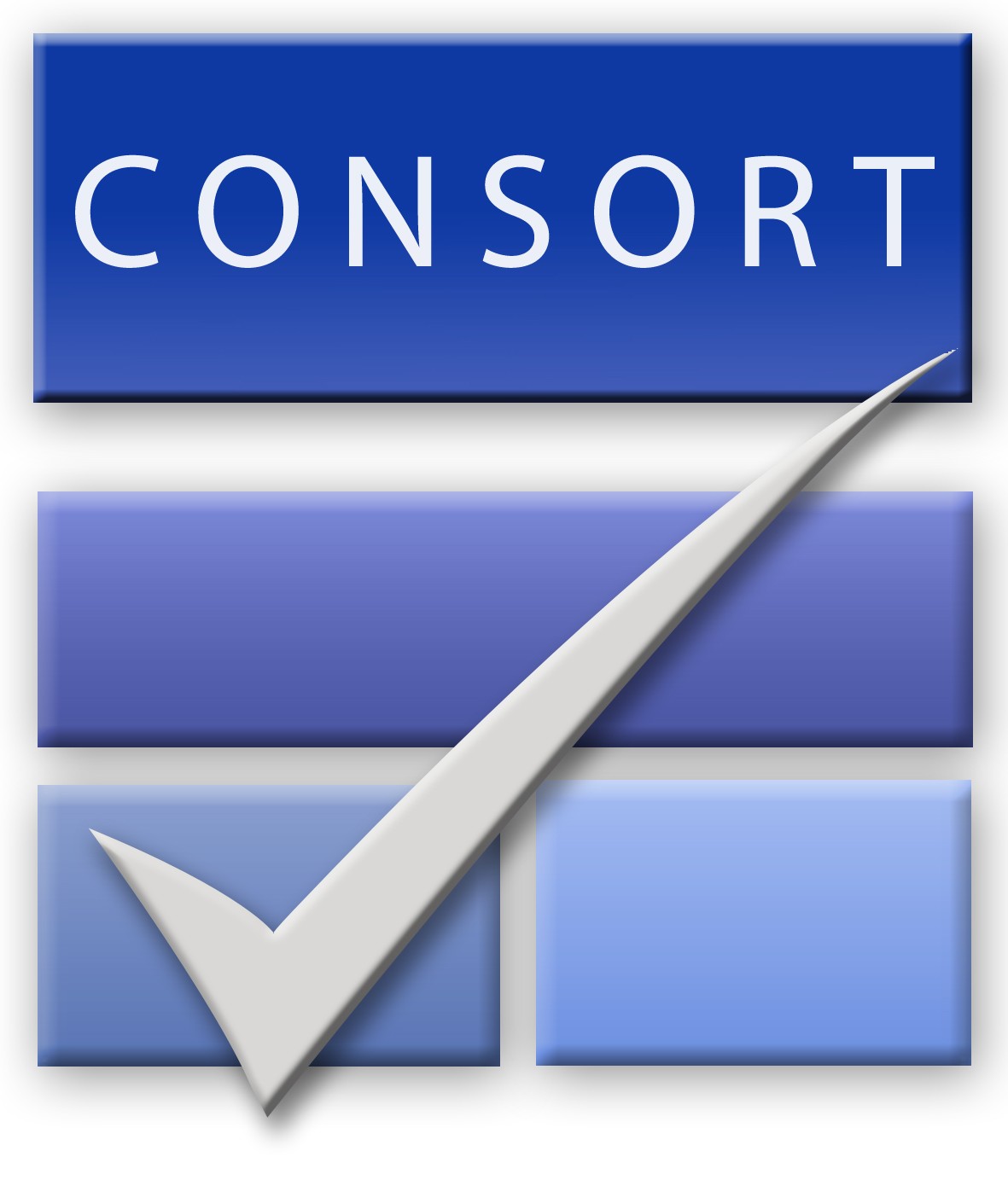
CONSORT 2010 checklist of information to include when reporting a randomised trial*

| Section/Topic | Item No | Checklist item | Reported on page No |
| --- | --- | --- | --- |
| Title and abstract | | | |
|  | 1a | Identification as a randomised trial in the title | 1 |
|  | 1b | Structured summary of trial design, methods, results, and conclusions (for specific guidance see CONSORT for abstracts) | 2 |
| Introduction | | | |
| Background and objectives | 2a | Scientific background and explanation of rationale | 4 |
|  | 2b | Specific objectives or hypotheses | 4 |
| Methods | | | |
| Trial design | 3a | Description of trial design (such as parallel, factorial) including allocation ratio | 4-5 |
|  | 3b | Important changes to methods after trial commencement (such as eligibility criteria), with reasons | N/A |
| Participants | 4a | Eligibility criteria for participants | 5 |
|  | 4b | Settings and locations where the data were collected | 4-5 |
| Interventions | 5 | The interventions for each group with sufficient details to allow replication, including how and when they were actually administered | 5-6 |
| Outcomes | 6a | Completely defined pre-specified primary and secondary outcome measures, including how and when they were assessed | 6 |
|  | 6b | Any changes to trial outcomes after the trial commenced, with reasons | N/A |
| Sample size | 7a | How sample size was determined | 7 |
|  | 7b | When applicable, explanation of any interim analyses and stopping guidelines | N/A |
| Randomisation: |  |  |  |
| Sequence generation | 8a | Method used to generate the random allocation sequence | 5 |
|  | 8b | Type of randomisation; details of any restriction (such as blocking and block size) | 5 |
| Allocation concealment mechanism | 9 | Mechanism used to implement the random allocation sequence (such as sequentially numbered containers), describing any steps taken to conceal the sequence until interventions were assigned | 5 |
| Implementation | 10 | Who generated the random allocation sequence, who enrolled participants, and who assigned participants to interventions | 5 |
| Blinding | 11a | If done, who was blinded after assignment to interventions (for example, participants, care providers, those assessing outcomes) and how | 5-6 |
|  | 11b | If relevant, description of the similarity of interventions | 5 |
| Statistical methods | 12a | Statistical methods used to compare groups for primary and secondary outcomes | 7 |
|  | 12b | Methods for additional analyses, such as subgroup analyses and adjusted analyses | 7 |
| Results | | | |
| Participant flow (a diagram is strongly recommended) | 13a | For each group, the numbers of participants who were randomly assigned, received intended treatment, and were analysed for the primary outcome | Figure 2 |
|  | 13b | For each group, losses and exclusions after randomisation, together with reasons | N/A |
| Recruitment | 14a | Dates defining the periods of recruitment and follow-up | 5, 6 |
|  | 14b | Why the trial ended or was stopped | 5 |
| Baseline data | 15 | A table showing baseline demographic and clinical characteristics for each group | 14 |
| Numbers analysed | 16 | For each group, number of participants (denominator) included in each analysis and whether the analysis was by original assigned groups |  |
| Outcomes and estimation | 17a | For each primary and secondary outcome, results for each group, and the estimated effect size and its precision (such as 95% confidence interval) | 7,8 |
|  | 17b | For binary outcomes, presentation of both absolute and relative effect sizes is recommended | 8 |
| Ancillary analyses | 18 | Results of any other analyses performed, including subgroup analyses and adjusted analyses, distinguishing pre-specified from exploratory | 8 |
| Harms | 19 | All important harms or unintended effects in each group (for specific guidance see CONSORT for harms) | 8 |
| Discussion | | | |
| Limitations | 20 | Trial limitations, addressing sources of potential bias, imprecision, and, if relevant, multiplicity of analyses | 10 |
| Generalisability | 21 | Generalisability (external validity, applicability) of the trial findings | 10 |
| Interpretation | 22 | Interpretation consistent with results, balancing benefits and harms, and considering other relevant evidence | 9 |
| Other information | | |  |
| Registration | 23 | Registration number and name of trial registry | 3 |
| Protocol | 24 | Where the full trial protocol can be accessed, if available | 11 |
| Funding | 25 | Sources of funding and other support (such as supply of drugs), role of funders | 11 |

*We strongly recommend reading this statement in conjunction with the CONSORT 2010 Explanation and Elaboration for important clarifications on all the items. If relevant, we also recommend reading CONSORT extensions for cluster randomised trials, non-inferiority and equivalence trials, non-pharmacological treatments, herbal interventions, and pragmatic trials. Additional extensions are forthcoming: for those and for up to date references relevant to this checklist, see [www.consort-statement.org](http://www.consort-statement.org/).
