## Supplemental Material for "Improving Proteinuria Screening with Mailed Smartphone Urinalysis Testing in Previously Unscreened Patients with Hypertension: a Randomized Controlled Trial"

The smartphone urinalysis app (Dip.io) analyzes urinalysis results using patients’ smartphone cameras. The home screening intervention flow is outlined below: Consented patients received a text message link to download the dip.io app from the Apple Store or Google Play. The testing kit, along with a project leaflet, were shipped by Healthy.io using a third-party fulfillment service. If participants had not completed the test within the next week, Healthy.io’s call center contacted participants to validate receipt of kit and application and trouble-shoot any issues that may have prevented participants from completing the test.

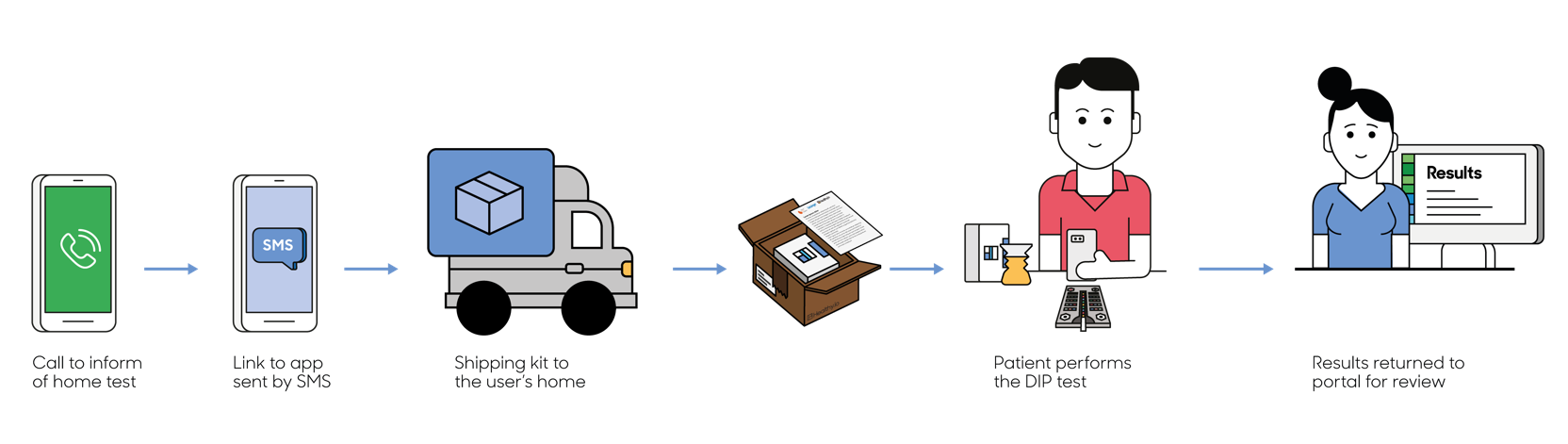

The Healthy.io DIP test consists of a home test kit and a smartphone application. The kit consists of a standard 10 parameter urinalysis dipstick, a custom designed urine cup and a color-board, which enables accurate analysis in different lighting environments. To conduct the test, patients open the app, follow directions provided on-screen, collect urine in the provided container, dip urinalysis dipstick (Acon Mission) and place it on the color board. They then scan the dipstick and color board using the app. Results are transmitted to a HIPAA-compliant web server in real-time and made available for physician review on the Healthy.io web portal or directly in the patient’s electronic health record.
